# A role for vitamin D and omega-3 fatty acids in major depression? An exploration using genomics

**DOI:** 10.1101/516013

**Authors:** Yuri Milaneschi, Wouter J Peyrot, Michel G Nivard, Hamdi Mbarek, Dorret I Boomsma, Brenda WJH Penninx

**Affiliations:** Department of Psychiatry, Amsterdam Public Health and Amsterdam Neuroscience, Amsterdam UMC, Vrije Universiteit/GGZ inGeest, Amsterdam, the Netherlands; Department of Biological Psychology, VU University Amsterdam, Amsterdam, the Netherlands; Qatar Genome Programme, Qatar Foundation, Doha, Qatar

**Keywords:** vitamin D, omega 3, major depression, genomics, etiology, causality

## Abstract

**Background:** Trials testing the effect of vitamin D or omega-3 fatty acids (n3-PUFA) supplementation on major depressive disorder (MDD) reported conflicting findings. These trials were boosted by epidemiological evidence suggesting an inverse association of circulating 25-hydroxyvitamin D (25-OH-D) and n3-PUFA levels with MDD. Observational associations may emerge from unresolved confounding, shared genetic risk, or direct causal relationships. We explored the nature of these associations exploiting data and statistical tools from genomics.

**Methods:** Results from GWAS on 25-OH-D (N = 79366), n3-PUFA (N = 24925) and MDD (135458 cases, 344901 controls) were applied to individual-level data (>2,000 subjects with measures of genotype, DSM-IV lifetime MDD diagnoses and circulating 25-OH-D and n3-PUFA) and summary-level data analyses. Shared genetic risk between traits was tested by polygenic risk scores (PRS). Two-sample Mendelian Randomization (2SMR) analyses tested the potential bidirectional causality between traits.

**Outcome:** In individual-level data, PRS were associated with the phenotype of the same trait (PRS 25-OH-D *p* = 1.4e-20, PRS N3-PUFA *p* = 9.3e-6, PRS MDD *p* = 1.4e-4), but not with the other phenotypes, suggesting a lack of shared genetic effects. In summary-level data, 2SMR analyses provided no evidence of a causal role on MDD of 25-OH-D (*p* = 0.50) or n3-PUFA (*p* = 0.16), or for a causal role of MDD on 25-OH-D (*p* = 0.25) or n3-PUFA (*p* = 0.66).

**Conclusions:** Applying genomics tools indicated that that shared genetic risk or direct causality between 25-OH-D, n3-PUFA and MDD is unlikely: unresolved confounding may explain the associations reported in observational studies. These findings represent a cautionary tale for testing supplementation of these compounds in preventing or treating MDD.

**Research in context:** *Evidence before this study:* Meta-analyses of trials testing the effect of vitamin D or omega-3 fatty acids (n3-PUFA) supplementation on major depressive disorder (MDD) reported conflicting findings, including small clinical effect or no effect. These trials were boosted by epidemiological evidence suggesting an inverse association of circulating 25-hydroxyvitamin D (25-OH-D) and n3-PUFA levels with MDD. However, observational associations may emerge from different scenarios, including unresolved confounding, shared genetic risk, or direct causal relationships.

*Added value of this study:* Genomics provides unique opportunities to investigate shared risk and causality between traits applying new statistical tools and results from genome-wide association studies (GWAS). In the present study we examined the nature of the association of 25-OH-D and n3-PUFA with MDD using the latest data and tools from genomics. We found no significant evidence of shared genetic risk or direct causality between vitamin D or n-3 PUFA and MDD; at this stage, unresolved confounding should be considered the most likely explanation for the association reported by observational studies.

*Implications of all the available evidence:* Findings from the present study, in conjunction with previous conflicting evidence from clinical studies, represent a cautionary tale for further research testing the potential therapeutic effect of vitamin D and n3-PUFA supplementation on depression, as the expectations of a direct causal effect of these compounds on mood should be substantially reconsidered. Genomic tools could be efficiently employed to examine the nature of observational associations emerging in epidemiology, providing some indications on the most promising associations to be prioritized in subsequent intervention studies.

## INTRODUCTION

In recent years a growing interest has emerged in the potential value of nutritional supplementation in mood disorders. Large-scale randomized controlled trials such as MoodFOOD^1^ or VITAL-DEP^2^ have been established to test the effect of supplementation in the prevention of depression. The rationale for these interventions stems from epidemiological evidence of an inverse association between depression and circulating concentrations of compounds that could be easily supplemented, such as vitamin D and omega-3 polyunsaturated fatty acids (n3-PUFA). Meta-analyses of large observational studies consistently reported cross-sectional, and to a lesser extent longitudinal, associations of 25-hydroxyvitamin-D (25-OH-D, the body reserve of vitamin D) and n3-PUFA with depression.^3,4^ In ∼2,500 psychiatrically well-characterized participants from the Netherlands Study of Depression and Anxiety, we previously confirmed^5,6^ lower concentrations of 25-OH-D and n3-PUFA in patients with depression as compared to healthy controls, and inverse associations with symptom severity. Previous clinical studies attempted to translate this observational evidence in interventions: meta-analyses^7–10^ pooling results from trials testing the effect on depression of supplementation vitamin D in up to ∼5000 cases and omega-3 in up to ∼1200 cases reported conflicting findings, with studies showing both small clinical effect or no effect. Conflicting findings have been commonly attributed to heterogeneity and limitations in study design, including small samples, inadequacy of supplementation dose or lack of blood markers to trace the biological availability of the compounds supplemented. The implementation of larger and methodologically robust trials has been repeatedly advocated to overcome this impasse.

Before moving to further intervention studies it is however crucial to obtain a clearer knowledge of the exact nature of an observational association, which could emerge from different scenarios. In the first of these scenarios observational associations emerging in nutritional epidemiology may be the product of unresolved confounding, as recently highlighted by Ioannidis^11^, since almost all nutritional variables are correlated with one another and with many social and behavioral factors. In the second scenario, an observational association may also be detected when both traits are epiphenomena stemming from the same etiological root, sharing part of their genetic liability. In the third scenario, the most favorable for for clinical translation, an observational association is detected because a trait (25-OH-D/n3-PUFA) is effectively causal for another (MDD). For vitamin D, hypotheses based on preclinical data suggested a potential impact of vitamin D on brain structure (vitamin D receptor is expressed in prefrontal cortex, amygdala and hippocampus) and pathophysiological processes relevant for mood.^12^ Similarly, n3-PUFA may exert important biological effects in the nervous system, modulating membrane fluidity, serotonergic transmission and inflammatory response^13^. Finally, the fourth scenario involves reverse causation, since depression may impact on 25-OH-D and n3-PUFA levels via habits related for instance to sunlight exposure or diet.^14^

It is not trivial to disentangle from which of the above scenarios an observational correlation arises. Genetic research has made enormous progress over the last years, and now provides unique opportunities to investigate shared risk and causality between traits applying new statistical tools and results from genome-wide association studies (GWAS). In the present study, we leveraged on results from the largest GWAS on 25-OH-D (79,366 samples)^15^, n3-PUFA (24,925 samples)^16^ and MDD (135,458 cases and 344,901 controls)^17^. We estimated the degree of genetic overlap between traits using both individual-level data (in NESDA) and summary-level data analyses combining GWAS summary statistics. Using summary data, Mendelian randomization analyses explored the potential bidirectional causality between 25-OH-D, n3-PUFA and MDD.

## METHODS

### Target sample for individual-level data analyses

Individual-level data analyses were based on 2047 unrelated participants (67.2% females) of European ancestry from the Netherlands Study of Depression and Anxiety (NESDA). Detailed descriptions of the rationale, design and methods for the study are given elsewhere.^5,6^ Briefly, NESDA is an ongoing cohort study into the long-term course and consequences of depressive and anxiety disorders. In 2004–2007, 2981 participants aged 18– 65 years were recruited from the community (19%), general practice (54%) and secondary mental health care (27%) and were followed-up during biannual assessments. The research protocol was approved by the ethical committee of participating universities, and all respondents provided written informed consent.

Genotyping, quality control and imputation were previously described in details^18^ (see also supplemental materials s1). Genotype data were used to build a relationship matrix measuring genetic similarity, which was pruned (0.05 threshold, no closer relationships than second cousin) in order to identify unrelated participants.

As previously reported,^5,6,18^ presence of DSM-IV lifetime diagnosis of MDD was assessed using the Composite Interview Diagnostic Instrument (CIDI version 2.1, WHO) administered by specially trained research staff at baseline, 2-year and 4-year follow up. Healthy controls included participants without any lifetime psychiatric disorder. Depressive symptoms severity was measured (baseline, 1-year, 2-year and 4-year follow-up) using the 30-item self-report Inventory of Depressive Symptoms (IDS-SR_30_).^19^ For the present analyses, scores across assessments were averaged in order to index the participant’s stable exposure to depressive symptoms.

Concentrations of 25-OH-D (N = 2013) and n3-PUFA (N = 2010) were quantified in blood samples taken at baseline as previously described.^5,6^ Serum 25-OH-D was measured using isotope dilution—online solid phase extraction liquid chromatography–tandem mass spectrometry. N3-PUFA was included in a metabolite panel measured using a high-throughput proton Nuclear Magnetic Resonance (NMR) platform (Nightingale Health Ltd., Helsinki, Finland).

Differences in main characteristics between participants with MDD and healthy controls were analyzed with chi-square or t-test, as appropriate. The sex-adjusted association of 25-OH-D and n3-PUFA with MDD and IDS-SR_30_scores was estimated using logistic and linear regression models. All analyses were performed in R v.3.5.2 (R Project for Statistical Computing)

### GWAS summary statistics

Summary statistics were obtained from large GWAS of international consortia. The SUNLIGHT consortium^15^ performed a GWAS on circulating concentrations of 25-OH-D from in 79,366 samples. Summary statistics for N3-PUFA were obtained from the MAGNETIC NMR GWAS^16^ examining the same metabolomics platform adopted in NESDA on up to 24,925 individuals. The Psychiatric Genomics Consortium (PGC)^17^ performed an overarching meta-analysis^2^ of all available GWAS datasets with MDD, including 135,458 cases and 344,901 controls.

### Polygenic risk scores

A polygenic risk score (PRS) for MDD was built based on the full polygenic signal from the MDD GWAS (see supplemental materials s3.1 for details of SNP selection for all PRS). Since NESDA data were part of the MDD GWAS, we re-ran the final meta-analyses after removal of overlapping datasets. PRS were built according to LDpred method^20^, which has shown an improved predictive performance compared with other methods by modeling a prior on effect sizes and LD information. The fraction of causal SNPs was set at 5% consistently with the estimate for schizophrenia.^21^

The genetic architecture of 25-OH-D and n3-PUFA may be characterized by few biologically relevant loci with relatively larger effect size. Therefore, PRS for these traits were based on genome-wide significant loci, harboring relevant genes for the synthesis and metabolism of these markers. For 25-OH-D we selected the six independent SNPs reported in the discovery GWAS^15^, related to key genes such as GC (Vitamin D binding protein, transporting vitamin D to target tissue), DHCR7 (7-dehydrocholesterol reductase, involved in vitamin D synthesis) and CYP2R1 and CYP24A1 (both Cytrochrome P450 family enzymes, catalyzing the transformation of vitamin D in its active form). For N-3 PUFAs, we identified 7 independent genome-wide significant SNPs by processing GWAS summary statistics (supplemental materials s2). The top SNP was located in the 3’-UTR region of FADS1 (fatty acid desaturase enzyme, involved in fatty acids metabolism), confirming the biological relevance of the top GWAS hits for this trait. Consistently, other significant loci included relevant genes for hepatic lipid metabolisms, such as LIPC and GCKR. PRS for 25-OH-D and n3-PUFA were therefore calculated as the number of risk alleles weighted by their effect sizes from the discovery statistics.

All PRS were standardized to aid comparison and interpretation of the results.

### PRS analyses

Same- and cross-trait associations of the different PRS with 25-OH-D and n3-PUFA concentrations and with MDD diagnosis were estimated using regression models (linear for 25-OH-D and N3-PUFA and binary logistic for MDD) adjusted for sex and 10 ancestry-informative genetic principal components. The proportion of phenotypic variance explained by PRS was additionally estimated (supplemental materials s3.2). In additional analyses focusing on MDD cases, the association between the PRS and symptom severity measured by IDS-SR_30_ was also estimated adjusting for sex and principal components. All analyses were performed in R.

### Estimation of genome-wide genetic correlation

LD-score regression^22^ (LDSC) was applied to GWAS summary statistics in order to estimate the genetic covariance between traits captured by all genotyped SNPs using the dedicated software. In univariate LDSC, the slope obtained by regressing GWAS test statistics on SNP LD-score (the sum of linkage disequilibrium r^2^ with all other SNPs) provides an estimate of a trait SNP-heritability (*h^2^_SNP_* the total variance in liability explained by the joint effect of all genotyped SNPs). *h^2^_SNP_* estimates for MDD were transformed to the liability scale based on a lifetime risk of 0.15. In the bivariate LDSC extension, the slope obtained by regressing the products of the test statistics of two traits on the LD-scores provides an estimate of the genetic correlation (*rg*) between the two traits.

### Mendelian randomization

Two-sample Mendelian randomization (2SMR) analyses^23^ based on GWAS summary statistics were performed to test the potential causal role of 25-OH-D and n3-PUFA on MDD risk and, inversely, of MDD on 25-OH-D and n3-PUFA levels. MR infers causality by using sets of SNPs reliably associated with an exposure as instrument for this exposure and regressing SNP-exposure effects against SNP-outcome effects. For each trait used as exposure, genome-wide significant independent SNPs were selected as instruments (supplemental materials s4.1). The use of 2SMR allows to significantly increase the statistical power by leveraging on effect sizes estimated in large GWAS. 2SMR analyses were based on inverse-variance weighted fixed effects-meta-analysis, pooling test statistics of all SNPs, inversely weighted by their standard error. Main sensitivity analyses were based on weighted-median and weighted-mode causal estimators, providing consistent causal estimates even when the majority of instruments are violating MR assumptions. Appropriate analytical checks of heterogeneity were additionally performed, including Cochran’s Q test, single SNP and leave-one-out SNP analyses. Analyses were conducted using the MR-Base R package.^23^

## RESULTS

### Individual-level data analyses: descriptives

The NESDA dataset included 1700 participants with a diagnosis of lifetime MDD and 347 healthy controls. MDD cases, as compared to controls, were more likely to be females and had lower blood levels of 25-OH-D (Cohen’s *d* −0.25) and n3-PUFA (Cohen’s *d* −0.15) (Table 1). The odds of lifetime MDD was reduced by ∼ 20% for each SD increase in 25-OH-D levels (sex-adjusted OR = 0.78, 95%CIs = 0.70-0.87, *p* = 1.4e-5) and by ∼ 10% for each SD increase in n3-PUFA levels (sex-adjusted OR = 0.87, 95%CIs = 0.78-0.97, *p* = 1.1e-2). Sex-adjusted levels of 25-OH-D and n3-PUFA were associated with depressive symptoms severity (Figure 1). Levels of 25-OH-D were inversely and linearly related to IDS-SR_30_ scores, while the association between n3-PUFA and symptom severity was best approximated by a quadratic form indicating higher depressive symptoms especially below a certain level of n3-PUFA.

**Figure 1.**
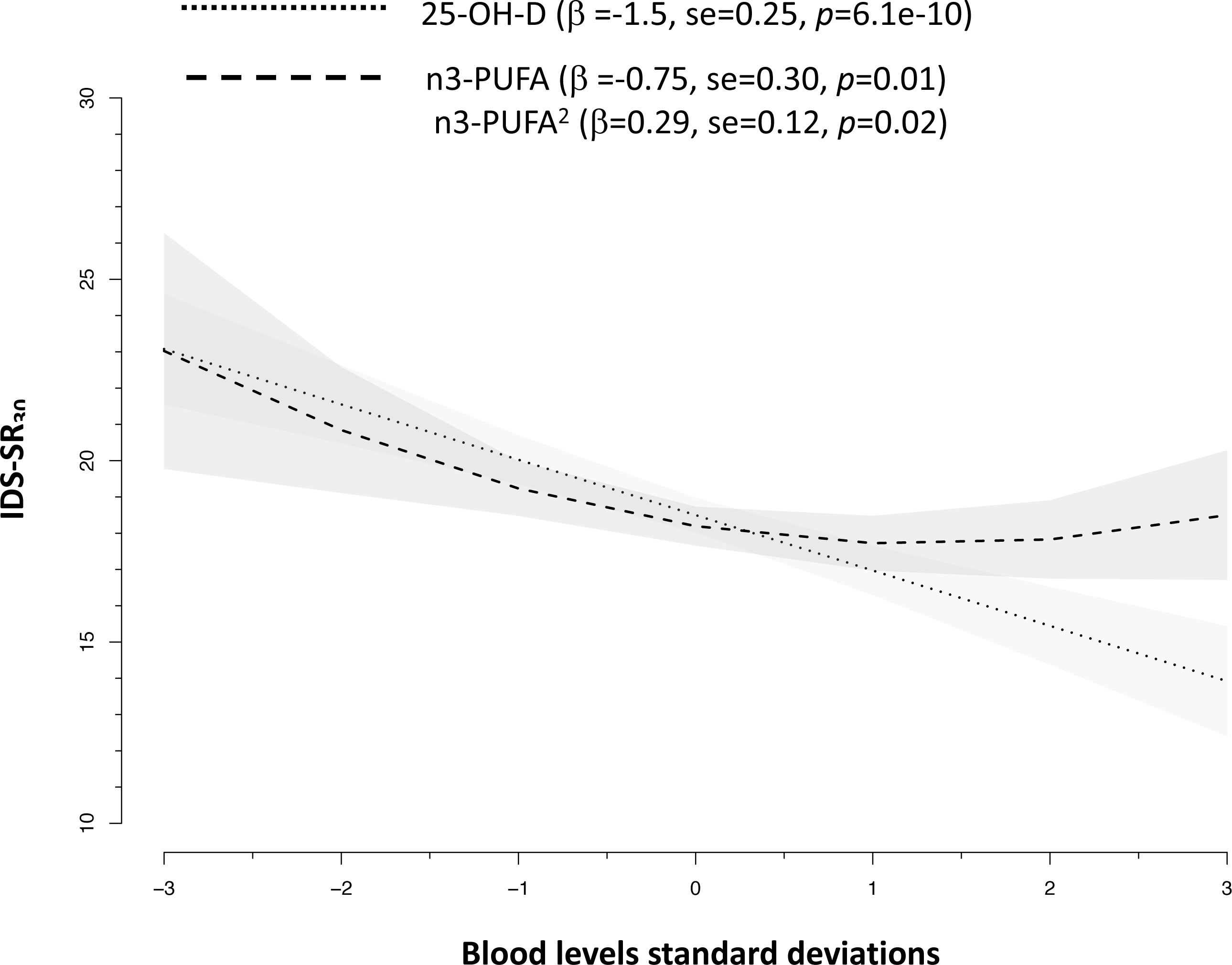
Associations between blood levels of 25-OH-D and n3-PUFA with depressive symptoms severity in 2047 NESDA participants. Slopes are adjusted for sex.

**Table 1.** Characteristics of the target sample (NESDA) for individual-level data analyses

| caracterisctis | MDD cases (*N* = 1700) | Controls (*N* = 347) | *P* |
| --- | --- | --- | --- |
| **Age** years (*mean* ± *SD*) | 42.4 (12.5) | 43.5 (14.2) | 0.17 |
| **Sex (F)** (%) | 68.6 | 60.1 | 4.3E-03 |
| **average IDS-SR<sub>30</sub>** (*mean* ± *SD*) | 21.7 (11.5) | 6.2 (5.0) | 2.6E-225 |
| **25-OH-D** (nmol/L) (*mean* ± *SD*) | 63.7 (28.3) | 70.5 (26.7) | 4.8E-05 |
| **N3-PUFAs** (nmol/L) (*mean* ± *SD*) | 0.38 (0.13) | 0.40 (0.14) | 1.6E-02 |

Unit of measure 25-OH-D, n3-PUFA: nmol/l (multiply by 0.4 to obtain ng/ml).

### Individual-level data analyses: PRS analyses

Table 2 shows same- and cross-trait associations of the PRS with 25-OH-D and N3-PUFA concentrations and with MDD. The association between PRS and phenotypes of the same trait were highly significant (PRS 25-OH-D *p* = 1.4e-20, PRS N3-PUFA *p* = 9.3e-6, PRS MDD *p* = 1.4e-4). The respective PRS explained 3.5% of trait variance for 25-OH-D level, 0.8% for N3-PUFA levels and 1.5% of MDD liability variance. Nevertheless, no association was found between PRS of one trait and the other phenotypes, suggesting a lack of shared genetic effects across traits.

**Table 2.** Same- and cross-trait associations of polygenic risk scores with circulating 25-hydroxyvitamin D, omega-3 fatty acids and Major Depressive Disorder in >2,000 participants from NESDA.

| PRS | 25-OH-D |  |  | N3-PUFA |  |  | MDD |  |  |
| --- | --- | --- | --- | --- | --- | --- | --- | --- | --- |
|  | N=2013 | mean = 64.8 (28.2) nmol/L |  | N=2010 | mean = 0.39 (0.13) nmol/L |  | 1700 cases + 347 controls |  |  |
| | $\beta$ | 95%CI | *p* | $\beta$ | 95%CI | *p* | OR | 95%CI | *p* |
| 25-OH-D | 5.29 | 4.18 - 6.39 | 1.4E-20 | -0.001 | -0.01 - 0.004 | 0.74 | 0.94 | 8.40 - 1.06 | 0.34 |
| n3-PUFA | -0.49 | -1.61 - 0.63 | 0.39 | 0.01 | 0.007 - 0.02 | 9.3E-06 | 1.01 | 0.90 - 1.14 | 0.83 |
| MDD | -0.57 | -1.70 - 0.55 | 0.32 | 0.001 | -0.03 - 0.01 | 0.58 | 1.26 | 1.12 - 1.41 | 1.4E-04 |

Results from linear (outcome: 25-OH-D, n3-PUFA) and binary logistic (outcome: MDD) regression analyses adjusted for sex and ten ancestry-informative genetic principal components.
Unit of measure 25-OH-D, n3-PUFA: nmol/l (multiply by 0.4 to obtain ng/ml).

**Table 2.** Two-samples Mendelian Randomization analyses based on GWAS summary statistics estimating causal effects between circulating 25-hydroxyvitamin D, omega-3 fatty acids and Major Depressive Disorder.

| Exposure | Outcome | N<br>SNPs | Inverse Variance<br>Weighted |  |  | Weighted Median |  |  | Weighted Mode |  |  |
| --- | --- | --- | --- | --- | --- | --- | --- | --- | --- | --- | --- |
|  |  |  | estimate | SE | *p* | estimate | SE | *p* | estimate | SE | *p* |
| 25-OH-D | MDD | 6 | -0.03 | 0.05 | 0.50 | -0.02 | 0.06 | 0.66 | -0.02 | 0.06 | 0.73 |
| n3-PUFA | MDD | 7 | -0.06 | 0.04 | 0.16 | -0.05 | 0.04 | 0.13 | -0.08 | 0.04 | 0.12 |
| MDD | 25-OH-D | 37 | -0.02 | 0.01 | 0.25 | -0.002 | 0.02 | 0.90 | 0.01 | 0.03 | 0.66 |
| MDD | n3-PUFA | 44 | 0.03 | 0.07 | 0.66 | 0.02 | 0.09 | 0.82 | -0.08 | 0.20 | 0.71 |

Units of measure in discovery GWAS:
- 25-OH-D, 1 unit increase in (log) concentrations; - n3-PUFA, 1 unit increase in (inverse normal transformation) concentrations; - MDD, 1 log-unit increase in risk.
SNP effects heterogeneity assessment in IVW:
- 25-OH-D causal for MDD, Cochran's *Q* $p=0.79$ ; - n3-PUFA causal for 25-OH-D, Cochran's *Q* $p=1.5e-3$ ; - MDD causal for 25-OH-D, Cochran's *Q* $p=0.15$ ; - MDD causal for n3-PUFA, Cochran's *Q* $p=0.17$

Consistently with these results, analyses focusing on lifetime MDD cases showed that only the PRS for MDD was associated with higher average 4-year severity of depressive symptom, explaining 0.7% of their variance (Figure 2). PRS for 25-OH-D and N3-PUFA were not related to symptoms severity in MDD cases.

**Figure 2.**
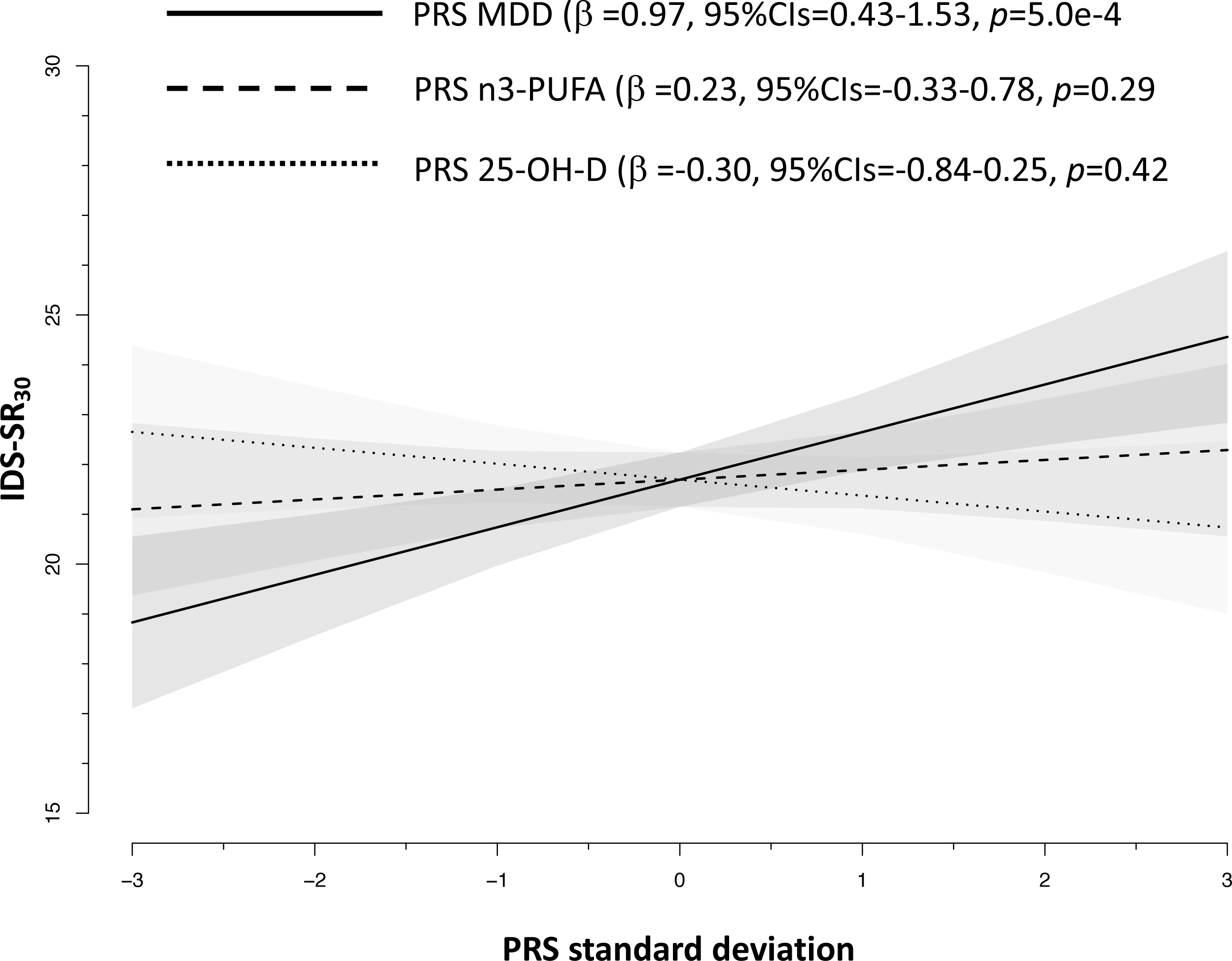
Associations between polygenic risk scores and 4-year average severity of depressive symptoms in 1700 cases with Major Depressive Disorder from NESDA. Slopes are adjusted for sex and 10 ancestry-informative principal components.

In additional analyses, we recalculated the PRS for 25-OH-D and n3-PUFA using the full polygenic signal of the discovery GWAS^15^ instead of selecting only genome-wide significant SNPs (supplemental materials s3.3). The resulting PRS were less strongly, although significantly, associated with the relative phenotype and explained a smaller proportion of variance (0.4% for 25-OH-D and 0.3% for N3-PUFA) as compared to the PRS based on the top SNPs, confirming that in the genetic architecture of 25-OH-D and N3-PUFA few biologically relevant loci may play a relative major role.

### Summary-level data analyses: genetic correlations

In univariate LDSC analyses, statistically significant *h^2^_SNP_* estimates were found for 25-OH-D (0.08, se = 0.02, *p* = 3.8e-5), N3-PUFA (0.15, se = 0.05, *p* = 1.4e-4) and MDD (0.08, se = 0.003, *p* = 1.3e-145). The estimates of *rg* between traits were small and non-significant (25-OH-D/MDD −0.06, se = 0.04, *p* = 0.12; n3-PUFA/MDD −0.06, se = 0.06, *p* = 0.35; 25-OH-D/n3-PUFA 0.15, se = 0.11, *p* = 0.18), confirming the absence of genome-wide shared genetic covariance between traits.

### Summary-level data analyses: Mendelian randomization

Table 3 reports the main results from 2SMR analyses. All approaches consistently indicated that the genetic instrument for 25-OH-D and n3-PUFA were not causally related to MDD risk. Of interest, only in the analyses focusing on n3-PUFA as exposure the heterogeneity test was significant (*p* = 1.5e-3), indicating the presence of at least one SNP potentially exhibiting horizontal pleiotropy. Inspection of the plots (eFigure 3) highlighted rs174546 (3’-UTR region of FADS1) as potential source of heterogeneity. In order to evaluate whether heterogeneity was indicating potential horizontal pleiotropy (the instrument/SNP is associated to the outcome through pathways others than the exposure), we performed a PheWAS (phenome-wide association scan, supplemental materials s4.3) confirming that the SNP was significantly associated with a wide array of traits beyond n3-PUFA, including for instance metabolic (e,g. cholesterol, triglycerides), cardiovascular (e.g. heart rate) or psychiatric (e.g. sleep duration, irritability) which are, in turn, associated with depression. In the opposite direction, 2SMR analyses examining MDD as exposure consistently showed no significant evidence of a causal role on 25-OH-D and n3-PUFA levels. Additionally, analyses using MR-Egger method^24^ were used and results are presented in eTable 2.

Although the number of instruments used to test the causal effect of 25-OH-D and n3-PUFA was lower than that (N = 10) recommended to run an adequately powered MR-Egger analyses, the estimates were substantially consistent with those obtained by other analyses, confirming the lack of potential causal effect of 25-OH-D and n3-PUFA on MDD and vice versa.

## DISCUSSION

In the present study we examined the nature of the association of 25-OH-D and n3-PUFA with MDD using the latest data and analytical tools developed in genomics. We found no significant evidence of shared genetic risk or direct causality between vitamin D or n-3 PUFA and MDD; unresolved confounding^11^ should be considered at this stage the most likely explanation for the association reported by observational studies.

Evidence of a common genetic base consistently shared between traits was not found. Even in data from the NESDA cohort, showing phenotypic associations of 25-OH-D and n-3 PUFA levels with MDD diagnosis presence and severity of depressive symptoms,^5,6^ the underlying polygenic risk for each trait was not associated with the phenotype of the others, indicating the lack of genetic overlap. Statistically non-significant genome-wide genetic correlations between 25-OH-D and MDD and n3-PUFA and MDD were previously reported.^15,17^ Nevertheless, those results may have been affected by the use of data for depression from GWAS relatively underpowered as compared to those available in the present study, and by different genetic architectures of depression (highly polygenic with many genetic variants with small effects) versus 25-OH-D or n3-PUFA (few genetic loci with relatively large effect sizes). In the present analyses, despite the use of polygenic scores for 25-OH-D or n3-PUFA built only on GWAS significant loci including biologically relevant genes for their synthesis and metabolism, no cross-trait association was found, suggesting that the biological pathways encompassing those genes are not shared with depression.

Furthermore, different MR analyses provided no evidence of a direct causal effect of 25-OH-D or n3-PUFA levels on MDD risk, despite the use of strong genetic instruments and the use of 2SMR techniques leveraging data from large GWAS studies. Furthermore, a potential causal effect of MDD on 25-OH-D or n-3PUFA levels was not detected, suggesting that also reverse causation does not explain the phenotypic association between these traits.

The present findings are consistent with other emerging evidence. A concomitant analysis^25^ applying MR techniques sustains the lack of causal effect of 25-OH-D on MDD, although the study was limited by the use of data for depression from a smaller GWAS (∼one third of the cases available here) and the lack of testing the potential reverse effect of MDD on 25-OH-D. Furthermore, a first study^26^ applying MR in a single (low powered) pregnancy cohort, found no evidence of a significant causal effect of n3-PUFA on antenatal, perinatal and postnatal depression.

Findings from the present study suggest that the observational associations vitamin D or n-3 PUFA and MDD commonly reported in epidemiological studies may be likely explained by unresolved confounding. The relationship of blood levels of 25-OH-D and n3-PUFA with MDD may indeed be confounded by several factors such health-related lifestyles (e.g. physical activity, diet), personality traits (e.g. conscientiousness and neuroticism) and comorbidity (e.g. obesity).^27,28^ Interestingly, we showed here that the top genetic variant linked to n3-PUFA levels on the FADS1 gene, is also associated with a wide array of traits, among which metabolic (e,g. cholesterol, triglycerides), cardiovascular (e.g. heart rate) and psychiatric (e.g. sleep duration, irritability) features that are, in turn, associated with MDD.^14^ Despite the fact that the relationship of MDD with 25-OH-D or n2-PUFA may be confounded and not be direct, epidemiological evidence consistently showing that depressed patients have lower blood level of these biological markers still raises an important concern for other health problems. For instance, we previously reported^5^ that one third of patients with established psychiatric diagnoses of depressive disorders had 25-OH-D blood concentrations considered insufficient and a risk for musculoskeletal complications. Normalization of vitamin D in depressed patients may therefore be important to prevent related complications and disability, in particular during late-life.

A major strength of the present study is the application of the latest techniques developed in genetic epidemiology, with analyses performed both at individual-level data, using information from a large and psychiatrically well-characterized cohort, and at summary-level data, combining GWAS summary statistics from the largest international consortia. An important limitation considering MR results is that genetic variants index the average lifetime exposure to risk factors and consequently could estimate an average lifetime causal effect without being able to identify a specific time in life or a specific portion of the risk factor distribution in which the causal effect may take place. Nevertheless, epidemiological data show that the association of 25-OH-D and n3-PUFA with MDD risk is consistent across different ranges of ages and that the functional form of these associations is linear without highlighting any consistent thresholds.^3–6^ Another major limitation is that both individual- and summary-level data analyses were based on samples of European ancestry and therefore results cannot be generalized to different populations.

Depression-related disability is a major health and economic burden for societies that require an urgent answer. Nevertheless, despite depression’s public health relevance, treatments options available nowadays are not optimal in terms of efficacy. Epidemiological evidence are constantly scanned in order to identify potentially actionable target that may substantially improve the prevention and treatment of depression. Recently developed genomic tools could be efficiently employed to examine the nature of observational associations emerging in epidemiology, providing some indications on the most promising associations to be prioritized in subsequent intervention studies. The premature and direct translation of observational associations in nutritional epidemiology into supplementation interventions may encounter indeed disappointing results, as emerging for different health outcomes. For instance, recent large-scale meta-analyses of randomized control studies have shown for instance that vitamin D supplementation does not prevent fractures, falls or improve bone mineral density,^29^ or that n3-PUFA supplementation had no significant effect on coronary heart disease or major vascular events.^30^ More recently, results from the large VITAL^31^ trial testing n3-PUFA supplementation in >25,000 participants showed no significant impact on the primary prevention of cardiovascular diseases or cancer.

In conclusion, we believe that findings from the present study, in conjunction with previous conflicting evidence from clinical studies, represent a cautionary tale for further research testing the potential therapeutic effect of vitamin D and n3-PUFA supplementation on depression, as the expectations of a direct causal effect of these compounds on mood should be substantially reconsidered. Upcoming results from already established large scale randomized controlled trials such as MoodFOOD^1^ or VITAL-DEP^2^ will tell whether this conclusion is correct.

## Acknowledgments

We would like to thank the investigators from the Psychiatric Genomics Consortium and the SUNLIGHT Consortium for sharing summary statistics of their GWAS.

In order to perform the present analyses we received summary statistics of the GWAS by Hyde et al. (Nat Genet. 2016; 48:1031-6) from 23andMe. We would like therefore to thank research participants and employees of 23andMe for making this work possible.

## Notes

**Funding:** For NESDA, funding was obtained from the Netherlands Organization for Scientific Research (Geestkracht program grant 10-000-1002); the Center for Medical Systems Biology (CSMB, NWO Genomics), Biobanking and Biomolecular Resources Research Infrastructure (BBMRI-NL), VU University’s Institutes for Health and Care Research (EMGO+) and Neuroscience Campus Amsterdam, University Medical Center Groningen, Leiden University Medical Center, National Institutes of Health (NIH, R01D0042157-01A, MH081802, Grand Opportunity grants 1RC2 MH089951 and 1RC2 MH089995). Part of the genotyping and analyses were funded by the Genetic Association Information Network (GAIN) of the Foundation for the National Institutes of Health.Computing was supported by BiG Grid, the Dutch e-Science Grid, which is financially supported by NWO.

