## supplemental materials for "A role for vitamin D and omega-3 fatty acids in major depression? An exploration using genomics"

***s1. Genotyping, quality control and imputation***	pag.2

***s2.Processing of n3-PUFA GWAS summary statistics***	pag.3

***eTable 1***	pag.3

***eFigure 1***	pag.3

***s3. PRS analyses***	pag.4

***s3.1 PRS generation***	pag.4

***s3.2 Estimation of variance explained by the PRS***	pag.4

***s3.3 PRS for 25-OH-D and n3-PUFA based on full polygenic signal***	pag.4

***s4. Mendelian randomization***	pag.6

***s4.1 Selection of independent SNPs as instruments for MR***	pag.6

***s4.2 MR plots***	pag.7

***eFigure 2***	pag.7

***eFigure 3***	pag.8

***eFigure 4***	pag.9

***eFigure 5***	*pag.10*

***s4.3 PheWAS for rs174546***	*pag.11*

***eFigure 6***	*pag.11*

***s4.4 MR-Egger***	*pag.12*

***eTable 2***	pag.12	***References***	pag.13

**URLs**	pag.16

***s1. Genotyping, quality control and imputation***

Genotyping, quality control and imputation were previously described in details^1^. Briefly, 95% of the samples were genotyped on the Affymetrix 6.0 Human SNP array and the remaining on the Perlegen-Affymetrix 5.0 array. After platform-specific QC the missing SNP genotypes between each platform were imputed using the GONL (Genome of the Netherlands)^2^ reference panel and then merged. Following more stringent QC, the SNPs from this cross-platform GONL imputed dataset (~1.2M) were used for a second round of imputations to the 1000G Phase 3 all ancestries reference panel using the Michigan Imputation Server^3^. Genotype data were used to build a relationship matrix measuring genetic similarity using GCTA^4^ , which was pruned (0.05 threshold, no closer relationships than second cousin) in order to identify unrelated participants.

***s2. Processing of n3-PUFA GWAS summary statistics***

Summary statistics for n3-PUFA were obtained from the MAGNETIC NMR GWAS^5^ examining the same metabolomics platform (Nightingale Health Ltd., Helsinki, Finland) adopted in NESDA on up to 24925 individuals. The original manuscript reports only global patterns of results across several metabolites. We processed the specific summary statistics for n3-PUFA in order to identify independent genome-wide significant SNPs by using the FUMA tool^6^. We applied the SNP2GENE function using GWAS summary statistics as an input and providing annotation of genomic areas identified by lead SNPs. The following criteria were applied: SNPs with MAF ≥ 0.01 were retained, the MHC region was excluded from annotation (due to long-range LD), an r^2^ 0.1 threshold was used to define LD structure of the lead SNP, maximum distance between LD blocks was set at 500kb and the EUR population of 1000Genomes was used as LD reference. Seven independent genome-wide significant SNPs from 5 genomic loci were identified.

**eTable 1** Independent genome-wide significant SNPs in n3-PUFA GWAS

| **rsid** | **chr** | **position** | ***p*** |
| --- | --- | --- | --- |
| rs174546 | 11 | 61569830 | 1.2E-34 |
| rs1260326 | 2 | 27730940 | 3.4E-14 |
| rs143988316 | 19 | 19667254 | 3.0E-12 |
| rs11604424 | 11 | 116651115 | 3.3E-10 |
| rs145717049 | 19 | 19130096 | 6.7E-09 |
| rs150617279 | 19 | 20139234 | 7.1E-09 |

**efigure 1** Manhattan plot for n3-PUFA GWAS

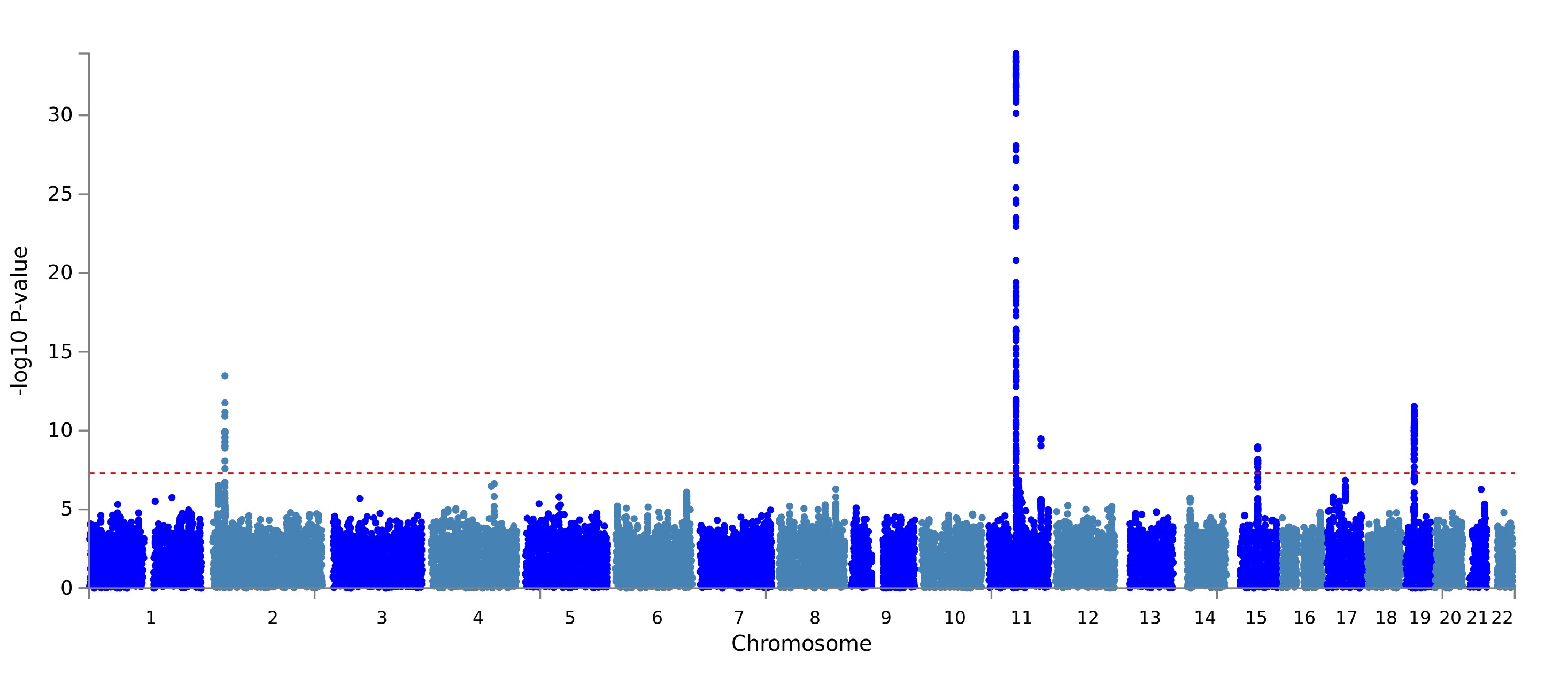

***s3. PRS analyses***

***s3.1 PRS generation***

PRS for MDD was built based on the full polygenic signal from the MDD GWAS.^7^ Since NESDA data were part of the GWAS, we re-ran the MDD meta-analyses after removal of overlapping datasets (~3K samples). Summary statistics of the discovery were filtered by removing In/Del and strand ambiguous variants, SNPs with INFO < 0.9, MAF < 0.01. Overlapping SNPs between the ~1.2M cross-platform GONL imputed and those retained from the discovery summary statistics were carried forward. PRS were built according to LDpred method^8^ using the dedicated software. 1000 unrelated individuals were selected to calculate LD for reference. The fraction of causal SNPs was set at 5% consistently with the estimate for schizophrenia by Palla and Dudbdrige.^9^

PRS for 25-OH-D and n3-PUFA were based on the independent SNPs associated at genome-wide significant level with the related traits in the discovery GWAS GWAS.^5,10^ In NESDA, the SNPs were selected in the 1000Genomes imputed set, see s1 section. The PRS for 25-OH-D and n3-PUFA were calculated as the number of risk alleles weighted by their effect sizes from the discovery statistics using PLINK.

***s3.2 Estimation of variance explained by the PRS***

Same- and cross-trait associations of the different PRS with 25-OH-D and n3-PUFA concentrations and with MDD diagnosis were estimated using regression models (linear for 25-OH-D and n3-PUFA and binary logistic for MDD) adjusted for sex and 10 ancestry-informative genetic principal components. In analyses with 25-OH-D or n3-PUFA as outcomes, the proportion of variance explained by the PRS was estimated based in the difference in R^2^ between a linear model including only covariates and a model additionally including the PRS. In analyses with MDD as outcome, Nagelkerke’s pseudo-R^2^ was derived and corrected for the covariates by substituting the null model in Nagelkerke’s equation for the model including the covariates. The corrected pseudo-R^2^ obtained was then re-scaled to the liability scale according to Lee et al.,^11^ obtaining a value directly comparable with heritability and robust against ascertainment bias. Linear transformation on the liability scale was based on lifetime risk (K) for MDD of 0.15.

***s3.3 PRS for 25-OH-D and n3-PUFA based on full polygenic signal***

In additional analyses, we recalculated the PRS for 25-OH-D and n3-PUFA using the full polygenic signal of the relative discovery GWAS instead of selecting only genome-wide significant SNPs. The new PRS was built using LDpred^8^ with the same protocol used for MDD PRS and assuming an infinitesimal model (all SNPs considered causal). The resulting PRS for 25-OH-D was associated with the phenotype of the same traits in NESDA (per SD increase, β =1.71, 95%CIs=0.59-2.84, *p*=2.9e-3), but explained a substantially smaller proportion of variance (0.4%) as compared to the PRS based on the top 6 SNPs (3.5%). Similarly, the PRS for n3-PUFA was associated with its circulating levels in NESDA (per SD increase, β = 0.01, 95%CIs= 0.002-0.01, *p*=9.9e-3), but explained a smaller proportion of variance (0.3%) as compared to the PRS based on the top 7 SNPs (0.8%). These results confirm that in the genetic architecture of 25-OH-D and n3-PUFA few biologically relevant loci may play a relative major role.

***s4. Mendelian randomization***

***s4.1 Selection of independent SNPs as instruments for MR***

Two-sample Mendelian randomization (2SMR) analyses^12^ based on GWAS summary statistics were performed to test the potential causal role of 25-OH-D and n3-PUFA on MDD risk and, inversely, of MDD on 25-OH-D and n3-PUFA levels. For each trait used as exposure, genome-wide significant independent SNPs were selected as instruments.

For analyses focusing on 25-OH-D as exposure, the 6 independent genome-wide significant SNPs reported in the discovery GWAS^10^ were used. For analyses focusing on n3-PUFA as exposure, the 7 independent genome-wide significant SNPs (see s.2 section) were selected. However, since rs145717049 was not present in summary statistics of MDD GWAS^7^ we replaced it with its best LD proxy (r^2^ = 0.3, from LDlink 3.3.0) in EUR population of 1000Genomes.

For MR analyses focusing on MDD as exposure and 25-OH-D as outcome, we selected 37 independent SNPs from the PGC MDD GWAS. In order to maximize the number of overlapping variants across the two GWAS with a different number of interrogated SNPs (due to difference in imputation reference panel: 1000 Genomes for MDD and HapMap2 for 25-OH-D) we performed the following selection steps: firstly, we identified 873 non strand-ambiguous SNPs (with *p* < 5.0e-8 in the MDD GWAS) available in both GWAS; we then retained only one SNP from the extended MHC region (due to long-range LD) and the rest was clumped using PLINK v1.9 using a 500kb window and an r^2^ 0.1 with the EUR population of 1000Genomes as LD reference.

We applied the same selection steps for MR analyses focusing on MDD as exposure and ne-PUFA as outcome and we selected 44 independent SNPs from the PGC MDD GWAS to be used as instruments.

***s4.2MR plots***

Panels: A. 2SMR estimates; B. Single SNP analyses ; C. Leave-one-out analyses

Units of measure in discovery GWAS: 25-OH-D, 1 unit increase in (log) concentrations; n3-PUFA, 1 unit increase in (inverse normal transformation) concentrations; MDD, 1 log-unit increase in risk.

***eFigure 2.*** 2SMR analyses estimating causal effects of 25-OH-D on MDD

Panels: A. 2SMR estimates; B. Single SNP analyses ; C. Leave-one-out analyses

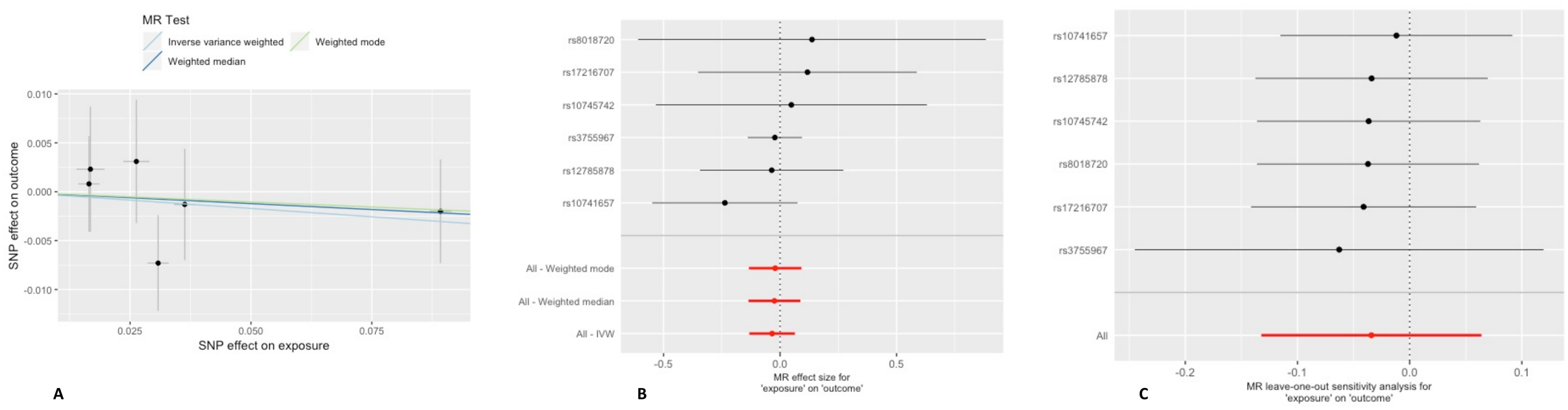

***eFigure 3.*** 2SMR analyses estimating causal effects of n3-PUFA on MDD

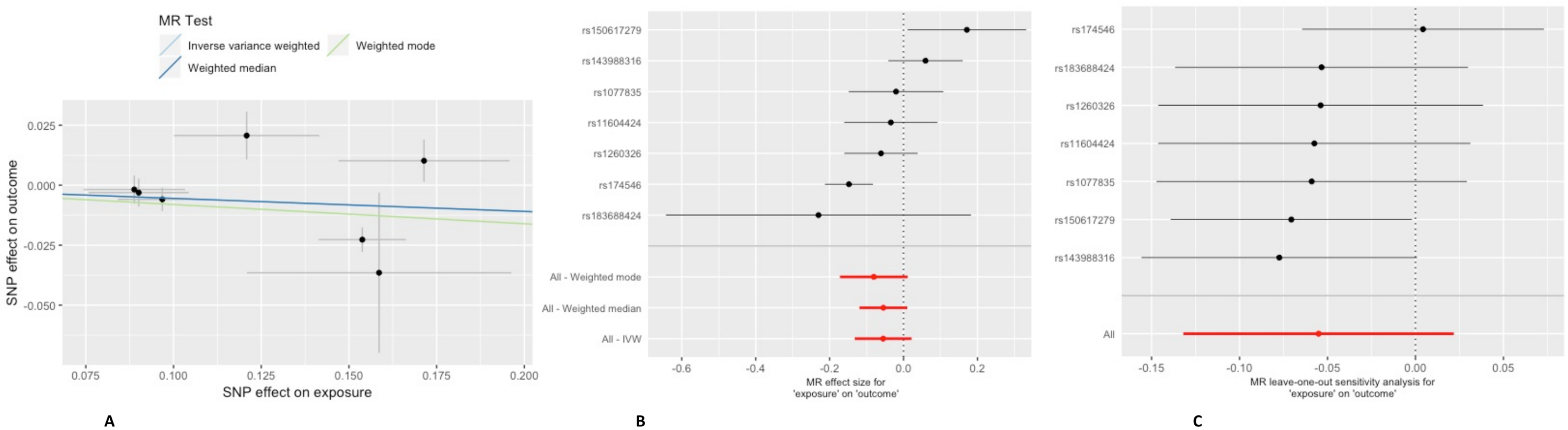

***eFigure 4.*** 2SMR analyses estimating causal effects of MDD on 25-OH-D

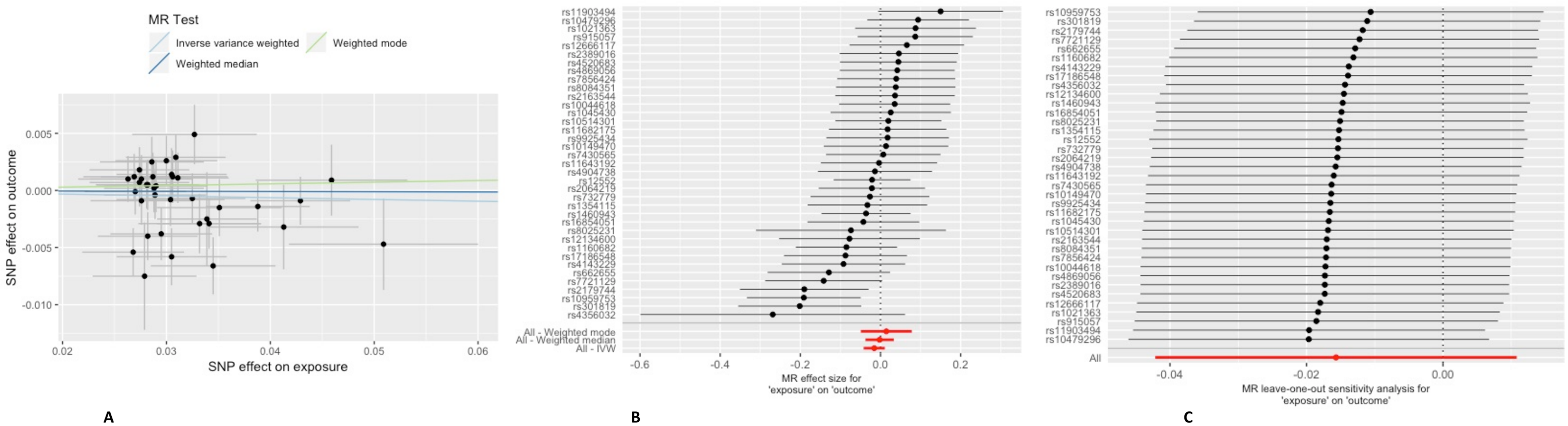

***eFigure 5.*** 2SMR analyses estimating causal effects of MDD on n3-PUFA

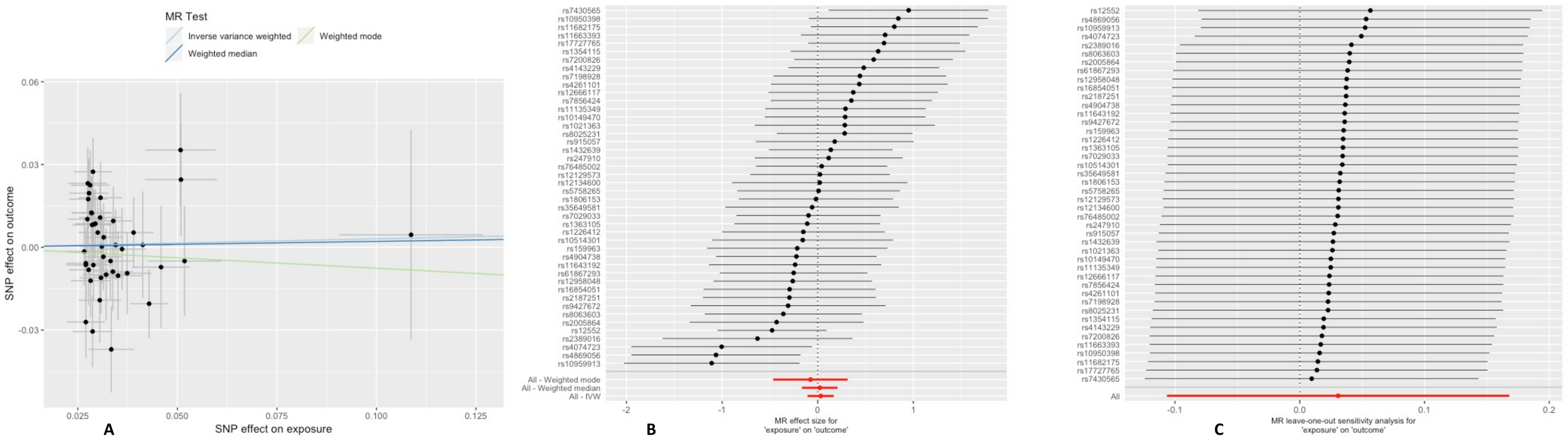

***s4.3PheWAS for rs174546***

In 2SMR analyses testing the causal effect of n3-PUFA on MDD we identified the SNP rs174546 as potential source of heterogeneity. In order to evaluate whether this heterogeneity was indicating potential horizontal pleiotropy (the instrument/SNP is associated to the outcome trough pathways others than the exposure), we performed a PheWAS (phenome-wide association scan) using the GWAS ATLAS Resource^14^. 3798 traits were scanned and 67 significant (0.05/3798 = 1.3e-05) associations were retrieved, including a wide array of traits.

***eFigure 6.*** Significant associations of rs174546 with traits organized in major domains

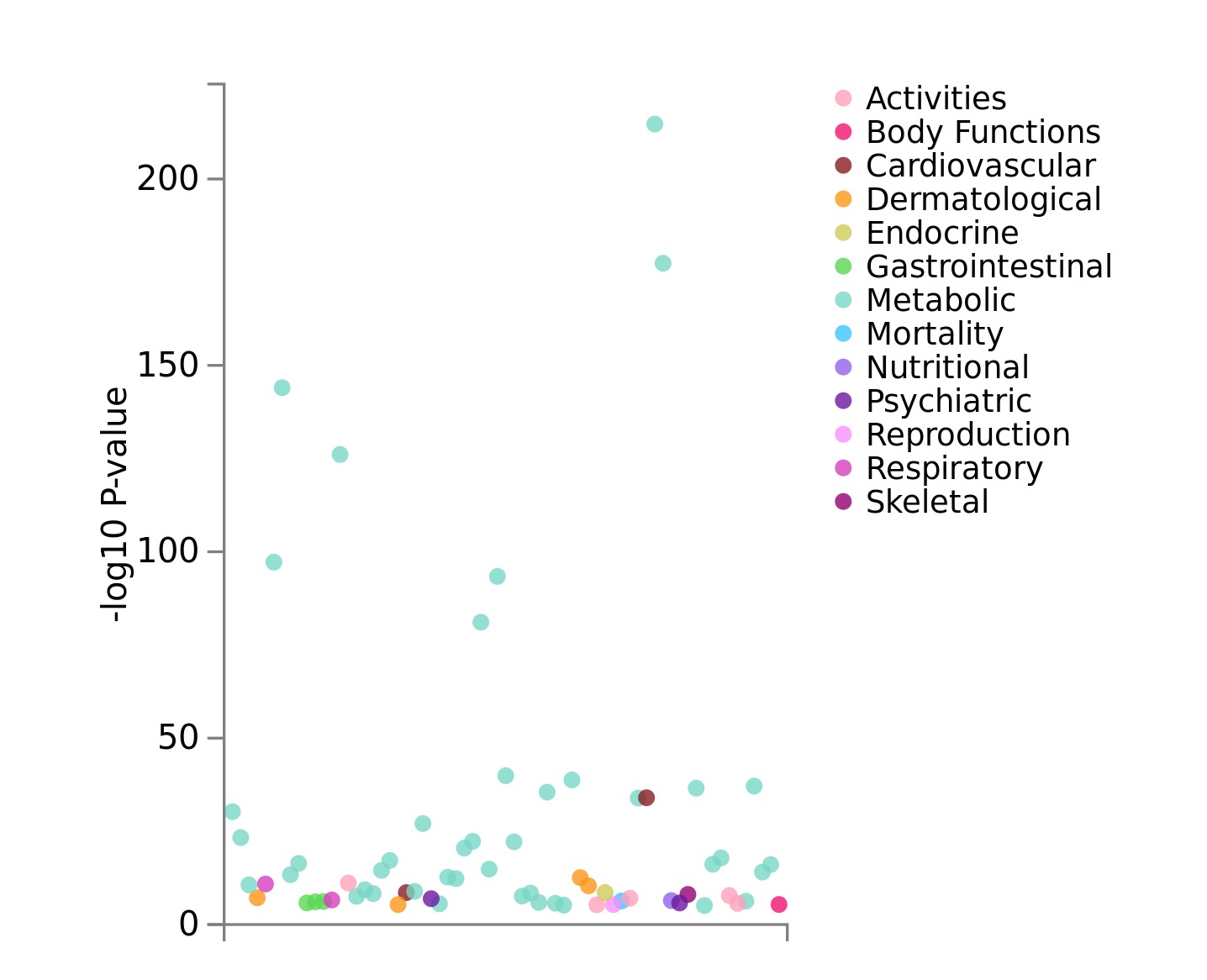

***s4.4MR-Egger***

Additional analyses were performed using the MR-Egger method^15^ relying on the InSIDE assumption (the magnitude of any pleiotropic effects should not correlate with the magnitude of the main effect) less conservative as the lack of horizontal pleiotropy The number of instruments used to test the causal effect of 25-OH-D and n3-PUFA was lower than that (N=10) recommended^15^ to run an adequately powered MR-Egger analyses. The estimates were substantially consistent with those obtained by other MR analyses. Furthermore, estimates from MR-Egger intercepts suggests the lack of substantial pleiotropy in analyses

***eTable 2*** 2SMR analyses using the MR-Egger method

|  |  |  |  | **MR-Egger** | | | | | | |
| --- | --- | --- | --- | --- | --- | --- | --- | --- | --- | --- |
|  |  |  |  | **Intercept** | | |  | **Slope** | | |
| **Exposure** | **Outcome** | **N SNPs** |  | **estimate** | **SE** | ***p*** |  | **estimate** | **SE** | ***p*** |
| **25-OH-D** | **MDD** | **6** |  | 0.004 | 0.01 | 0.74 |  | -0.03 | 0.09 | 0.71 |
| **n3-PUFA** | **MDD** | **7** |  | 0.01 | 0.02 | 0.71 |  | -0.12 | 0.16 | 0.50 |
| **MDD** | **25-OH-D** | **37** |  | 0.003 | 0.003 | 0.29 |  | -0.11 | 0.09 | 0.23 |
| **MDD** | **n3-PUFA** | **44** |  | 0.004 | 0.011 | 0.74 |  | -0.079 | 0.337 | 0.82 |

**URLs**

25-OH-D GWAS summary statistics <https://drive.google.com/drive/folders/0BzYDtCo_doHJRFRKR0ltZHZWZjQ>

MDD GWAS summary statistics <https://www.med.unc.edu/pgc/results-and-downloads> (the statistics publicly available are based on a GWAS not including 23andMe data, access to which is restricted by a Data Transfer Agreement)

LDSC <https://github.com/bulik/ldsc>

PLINK <https://www.cog-genomics.org/plink/1.9/>

LDpred <https://github.com/bvilhjal/ldpred>

MR-Base for two-samples MR <https://mrcieu.github.io/TwoSampleMR/>

mRnd, power calculator for MR <http://cnsgenomics.com/shiny/mRnd/>

FUMA <http://fuma.ctglab.nl/>

LDLink <https://ldlink.nci.nih.gov/>

ATLAS GWAS <http://atlas.ctglab.nl/>

CGTA <https://cnsgenomics.com/software/gcta/#Overview>
